# Zinc oxide nanoparticles modulate the gene expression of ZnT_1_ and ZIP_8_ to manipulate zinc homeostasis and stress-induced cytotoxicity in human neuroblastoma SH-SY5Y cells

**DOI:** 10.1101/2020.04.22.055152

**Authors:** Chien-Yuan Pan, Fang-Yu Lin, Lung-Sen Kao, Chien-Chang Huang, Pei-Shan Liu

## Abstract

Zinc ions (Zn^2+^) are important messenger molecules involved in various physiological functions. To maintain the homeostasis of cytosolic Zn^2+^ concentration ([Zn^2+^]_c_), Zrt/Irt-related proteins (ZIPs) and Zn^2+^ transporters (ZnTs) are the two families of proteins responsible for decreasing and increasing the [Zn^2+^]_c_, respectively, by fluxing Zn^2+^ across the membranes of the cell and intracellular compartments in opposite directions. Most studies focus on the cytotoxicity incurred by a high concentration of [Zn^2+^]_c_ and less investigate the [Zn^2+^]_c_ at physiological levels. Zinc oxide-nanoparticle (ZnO-NP) is blood brain barrier-permeable and elevates the [Zn^2+^]_c_ to different levels according to the concentrations of ZnO-NP applied. In this study, we mildly elevated the [Zn^2+^]_c_ by zinc oxide-nanoparticles (ZnO-NP) at concentrations below 1 μg/ml, which had little cytotoxicity, in cultured human neuroblastoma SH-SY5Y cells and characterized the importance of Zn^2+^ transporters in 6-hydroxy dopamine (6-OHDA)-induced cell death. The results show that ZnO-NP at low concentrations elevated the [Zn^2+^]_c_ transiently in 6 hr, then declined gradually to a basal level in 24 hr. Knocking down the expression levels of *ZnT*_*1*_ (mostly at the plasma membrane) and *ZIP*_*8*_ (present in endosomes and lysosomes) increased and decreased the ZnO-NP-induced elevation of [Zn^2+^]_c_, respectively. ZnO-NP treatment reduced the basal levels of reactive oxygen species and *Bax/Bcl-2* mRNA ratios; in addition, ZnO-NP decreased the 6-OHDA-induced ROS production, *p53* expression, and cell death. Therefore, mild elevations in [Zn^2+^]_c_ induced by ZnO-NP activate beneficial effects in reducing the 6-OHDA-induced cytotoxic effects. Therefore, brain-delivery of ZnO-NP can be regarded as a potential therapy for neurological disease.

## Introduction

Zinc ion (Zn^2+^) is essential for all living organisms and is the second most abundant transition element in human. It is a cofactor in many proteins regulating their catalytic activities and structure. In addition, recent emerging evidence has shown that Zn^2+^ is a messenger in regulation of many cellular activities such as cell cycle, cell proliferation, differentiation and death via different signaling pathways (1, 2). Cytosolic Zn^2+^ concentration ([Zn^2+^]_c_) changes during cell cycle, differentiation and cell death (3). During cell proliferation, the tyrosine phosphatases are suppressed by a small elevation of [Zn^2+^]_c_ to activate ERK pathway (4). A number of transcription factors, such as p53, contain Zn^2+^ binding motifs affecting cell cycle and survival (5).

The paradoxical, but vital roles of Zn^2+^ in nervous system have gained recognition recently (6, 7). Zn^2+^ is essential for neurogenesis, neuronal differentiation and synaptic transmission. The inhibition of synaptic Zn^2+^ signaling in hippocampus and amygdala by Zn^2+^ chelators affects cognition (8). Zn^2+^ deficiency reduces neurogenesis and associates with neuronal dysfunction. A correlation between Zn^2+^ deficiency and depression has been demonstrated in both clinical studies and animal models (9, 10). In contrast, high Zn^2+^ levels block mitochondrial function and induce apoptosis in the development of pathophysiology of CNS disorders including epilepsy, schizophrenia and Alzheimer’s Disease (11). At cellular level, high dose of Zn^2+^ is neurotoxic causing cell death (12–14) and Zn^2+^ deficiency causes caspase-dependent apoptosis in human neuronal precursor cells (15, 16). Zn^2+^ supplementation significantly reduces spinal cord ischemia-reperfusion injury in rats (17). However, dietary Zn^2+^ supplementation has restrictions and limitations in crossing brain-blood barrier (BBB), which has limited permeability for Zn^2+^, especially when the desired final Zn^2+^ level is higher than physiological levels (18). Thus, controlled and targeted delivery of Zn^2+^ is highly desirable.

Nanoparticles (NP) technologies have been used for the targeted delivery of chemicals (19). In nervous system, polylactide-co-glycolide or BBB ligand specific-modified polylactide polymers are used to carry Zn^2+^ across BBB (18, 19). However, the rate is slow, the cellular or brain entrance were evidenced after several days (19). We have previously demonstrated the entrance of zinc oxide-NP (ZnO-NP) into brain via olfactory bulb in rat and elevates the [Zn^2+^]_c_ in cultured cells (20). Therefore, ZnO-NP has the potential to be a potent means for Zn^2+^ delivery to regulate [Zn^2+^]_c_ homeostasis in the central nervous system.

The cellular uptake of ZnO-NP into intracellular compartments is via endocytosis followed by dissolution that occurs in acidic compartments to convert ZnO-NP to Zn^2+^ (20). Two classes of proteins are implicated in Zn^2+^ transport for [Zn^2+^]_c_ homeostasis: solute-linked carrier 30 (SLC30, Zn transporter (ZnT)) and SLC39 (Zrt/Irt-realted proteins (ZIP)) decrease and increase the [Zn^2+^]_c_, respectively, by fluxing Zn^2+^ across the membranes of cell and intracellular organelles in opposite directions. The ZIP proteins then transport the accumulated Zn^2+^ in these acidic compartments to the cytosol and ZnT proteins work corporately to flux Zn^2+^ out of the cytosol. Therefore, ZnO-NP may be different from direct Zn^2+^ application in regulating expression levels of Zn^2+^ transporters to control Zn^2+^ homeostasis.

ZnO-NP at high dosage causes apoptosis in lung (21) and neural stem cells (13) and interferes with the ion channel activities in primary cultured rat hippocampal neurons (22). However, toxicity is not seen under exposure to ZnO-NP at low doses, such as 6 ppm (70 μM)(13), or 10 μM (20). The importance of Zn^2+^ to normal functioning of the central nervous system is increasingly appreciated (9, 15). In this report, we mildly elevated the concentration of [Zn^2+^]_c_ in human neuroblastoma cells, SH-SY5Y, by ZnO-NP at concentrations below 1 μg/ml. ZnO-NP treatment greatly enhanced the expression level of ZnT_1_ and less affected the expression of ZIP_8_. ZnO-NP treatment decreased the basal level of reactive oxygen species (ROS) and the expression ratio of Bax/Bcl-2. In addition, ZnO-NP treatment recued the cell death caused by the 6-hydroxy dopamine (6-OHDA). Therefore, BBB-permeable ZnO-NP provides a therapeutic strategy to treat neurodegeneration disorders by fin-tuning the [Zn^2+^]_c_.

## Materials and Methods

### Chemicals

ZnO-NP were purchased from Sigma-Aldrich Co. (St. Louis, MO, USA). Their preparation followed protocols described in our previous work (21). The size range of ZnO-NP in solution was from 20 to 80 nm with an average of 45 nm. SH-SY5Y neuroblastoma cells were purchased from the American Type Culture Centre CRL2266 (Manassas, VA, USA). FluoZin-3-AM, reverse transcriptase III, and TRIzol^®^ reagent were purchased from Invitrogen Co. (Carlsbad, CA, USA). RNase-free DNAse I and RNeasy purification columns were purchased from Qiagen Inc. (Valencia, CA, USA). Random hexamer primers were obtained from Fermentas Inc. (Burlington, Canada). iQ SYBR Green Supermix was obtained from Bio-Rad Inc. (Hercules, CA, USA). Other chemicals were obtained from Merck KGaA (Darmstadt, German) otherwise indicated.

### Cell culture

Human neuroblastoma SH-SY5Y cells were cultured in minimal essential medium (Gibico 41500-034) supplemented with F12 nutrient mixture (Gibico 21700-075) and 10% fetal bovine serum. The cells were kept in a humidified 5%-CO_2_ incubator at 37 **°**C (20).

### [Zn^2+^]_c_ Measurements

Suspended cells were incubated in a Loading buffer (in mM, NaCl 150, glucose 5, Hepes 10, MgCl_2_ 1, KCl 5, CaCl_2_ 2.2, pH7.3) containing 10 μM of FluoZin-3-AM at 37°C for 30 minutes. After washing out the FluoZin-3-AM by centrifugation and resuspending the cell in Loading buffer, the changes in the fluorescence intensity were recorded as described before (20).

### RT-PCR assay

RNA extraction and reverse transcription were performed following the protocols suggested by the manufactures. The primers for the polymerase chain reactions (PCR, Q-Amp™ 2x HotStart PCR Master Mix) were listed in Supplementary Table S1. The products were separated by electrophoresis on 2% agarose gels, stained with ethidium bromide, and photographed with ultraviolet transillumination. For quantitative PCR (qPCR), the kit used was IQ^2^ Fast qPCR System and the instrument was from Illumina Inc. (Eco™ Real-time PCR system) (23).

### MTT assay

The MTT assay, an index of cell viability and cell growth, is based on the ability of viable cells to reduce MTT (3-(4,5-dimethylthiazol-2-yl)-2,5-diphenyl tetrazolium bromide) (24). All samples were assayed in triplicate, and the mean for each experiment was calculated. Five batches of cells were used in this experiments.

### ROS measurements

To quantify the production of ROS, we loaded the cells with 2’,7’-dichlorodihydrofluorescein diacetate (H_2_DCFDA, Molecular Probes®) and incubated at 37°C, 5% CO_2_ for 30 minutes. After replacing the medium, H_2_O_2_ or 6-OHDA were added. The fluorescence intensities were measured by a microplate reader (Glomax-multidetection system, Promega, USA) with excitation at 485 nm and emission at 500 - 560 nm.

### *ZIP*_*8*_ and *ZnT*_*1*_ shRNA knockdown

Plasmids expressing short hairpin RNAs (shRNA) against ZIP_8_ and ZnT_1_ were purchased from National RNAi Core Facility, Academia Sinica, Taiwan, and the target sequences of these shRNAs (4 for *ZIP*_*8*_ and 5 for *ZnT*_*1*_) were listed in Supplementary Table S2. Lipofectamine 2000^®^ (Invitrogen, Carlsbad, CA) was used to transfect these plasmids into SH-SY5Y cells(25). An apoplasmid was used as negative control.

### Statistical analysis

Statistical analysis was performed using one-way analysis of variance and significant differences were assessed by Student’s *t* test. A *p* value less than 0.05 was regarded as statistically significant.

## Results

### 1. ZnO-NP elevates [Zn^2+^]_c_ in cultured SH-SY5Y cells

To examine ZnO-NP at low doses can elevate [Zn^2+^]_c_ in cultured human neuroblastoma SH-SY5Y cells, we loaded the cells with FluoZin3, a Zn^2+^-sensitive dye, and monitored the changes in fluorescence intensities (Fig. 1). The addition of ZnO-NP (0.081 and 0.814 μg/ml) increased the fluorescence intensity gradually during the 200-s recording period in a concentration-dependent manner. For 25-hr long-term treatment, the fluorescence intensities measured reached a maximum in 6 hr when treated with different concentrations of ZnO-NP (0.081, 0.814, and 8.14 μg/ml). These results reveal that ZnO-NP apparently elevates the [Zn^2+^]_c_ transiently in a concentration- and time-dependent mode even at low concentrations.

**Figure 1.**
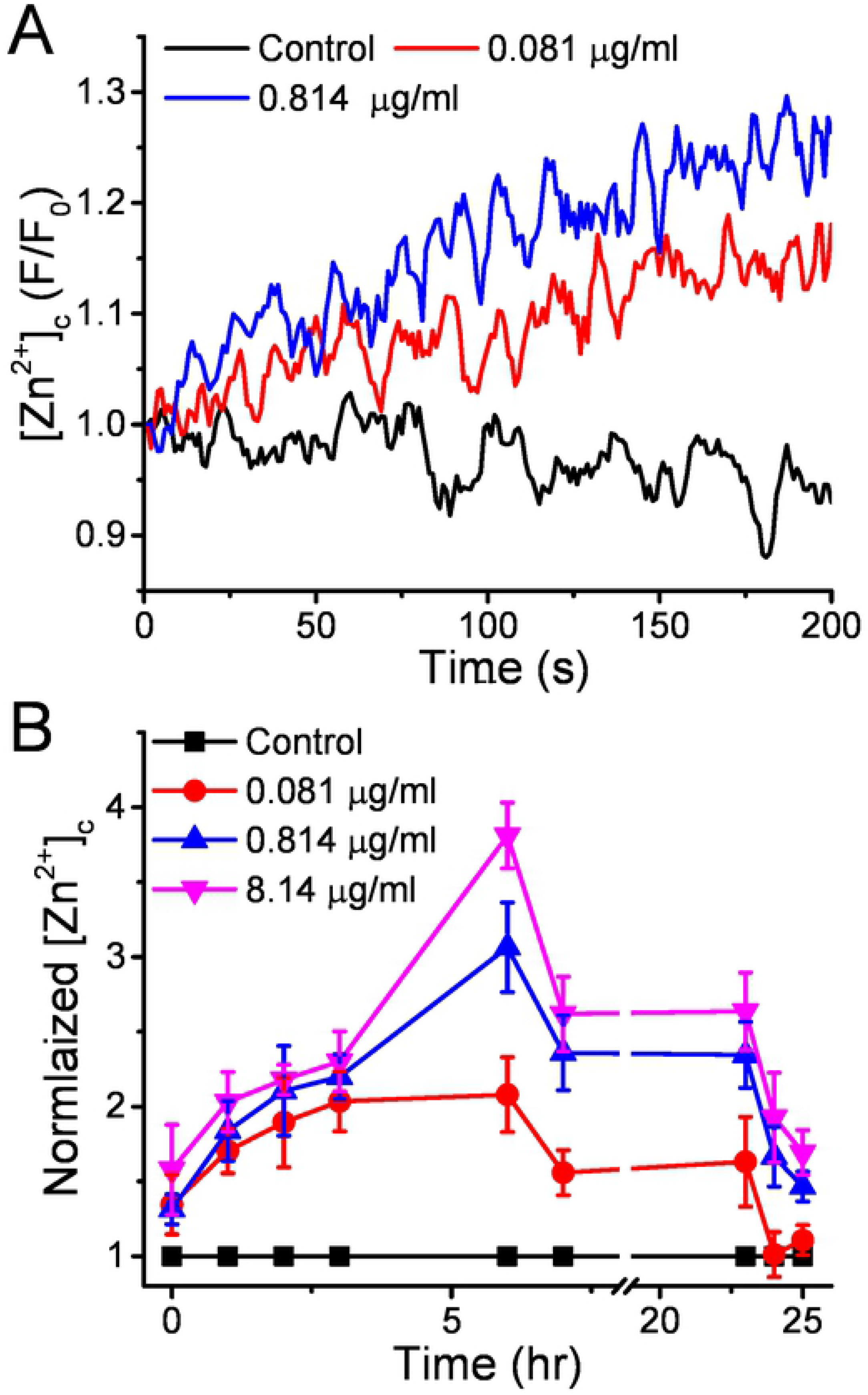
ZnO-NP exposure induces a transient elevation of [Zn^2+^]_c_ in SH-SY5Y cells. We loaded the cells with FluoZin-3 and monitored the changes of the fluorescence intensities from a group of suspended cells stimulated with different concentrations of ZnO-NPs. A. The short-term [Zn^2+^]_c_ responses. ZnO-NP (0, 0.081, and 0.814 μg/ml) were added at the beginning of the recording and the fluorescence intensities were normalized to the value at the time zero (F/F_0_). B. Long-term exposure of ZnO-NP. The fluorescence intensities from ZnO-NP-treated suspension cells were normalized to the control group without ZnO-NP treatment (Normalized [Zn^2+^]_c_) at different time after ZnO-NP exposure. Data presented were Mean ± SEM from 3 batches of cells.

### 2. ZnT_1_ and ZIP_8_ regulate the ZnO-NP-induced [Zn^2+^] responses in SH-SY5Y cells

ZIPs and ZnTs play important roles in maintaining the [Zn^2+^]_c_ homeostasis. We first characterized the expression levels of *ZnT* and *ZIP* isoforms in cultured SH-SY5Y cells by RT-PCR and the results showed significant expressions of *ZnT*_*1*_, *ZnT*_*3*_, *ZnT*_*4*_, *ZnT*_*5*_, *ZnT*_*6*_, *ZnT*_*7*_, *ZnT*_*9*_ and *ZnT*_*10*_ (Supplementary Fig. S1A) and *ZIP*_*1*_, *ZIP*_*3*_, *ZIP*_*4*_, *ZIP*_*6*_, *ZIP*_*7*_, *ZIP*_*8*_, *ZIP*_*9*_, *ZIP*_*10*_, *ZIP*_*11*_, *ZIP*_*13*_ and *ZIP*_*14*_ (Supplementary Fig. S1B). ZnT_1_ is the main transporter at the plasma membrane to efflux Zn^2+^ out of cells and lowers the [Zn^2+^]_c_ (26); ZIP_8_ presents in the synaptic vesicles and lysosomes to transport Zn^2+^ from intracellular compartments to the cytosol (27, 28). Since endocytosis is the main route for ZnO-NP entrance into the cell and dissolution into Zn^2+^ occurs in an acidic compartment (20), we focused on characterizing the involvement of ZnT_1_ and ZIP_8_ in modulating the ZnO-NP-induced [Zn^2+^]_c_ response in SH-SY5Y cells (Fig. 2). We adopted qPCR to investigate the mRNA levels of *ZnT*_*1*_ and *ZIP*_*8*_ in SH-SY5Y cells after the addition of ZnO-NP of different concentrations. The average results show that a low-dose of ZnO-NP (0.081 μg/ ml) elevated the expression levels of *ZnT*_*1*_ and *ZIP*_*8*_ transiently in 6 hr and then declined to a basal level after 24 hrs. High doses of ZnO-NP (0.814 and 8.14 μg/ ml) treatment maintained the expression of *ZnT*_*1*_ at a level 4~8 fold higher than the control group during the 24-hour exposure period. ZnO-NP at 0.814 μg/ml elevated and maintained the expression of *ZIP*_*8*_ at a level 2-3 fold higher than the control group, however, at 8.14 μg/ml, ZnO-NP had little effect on the expression of *ZIP*_*8*_. These results reveal that ZnO-NP exposure differentially enhances the expression of *ZnT*_*1*_ and *ZIP*_*8*_.

**Figure 2.**
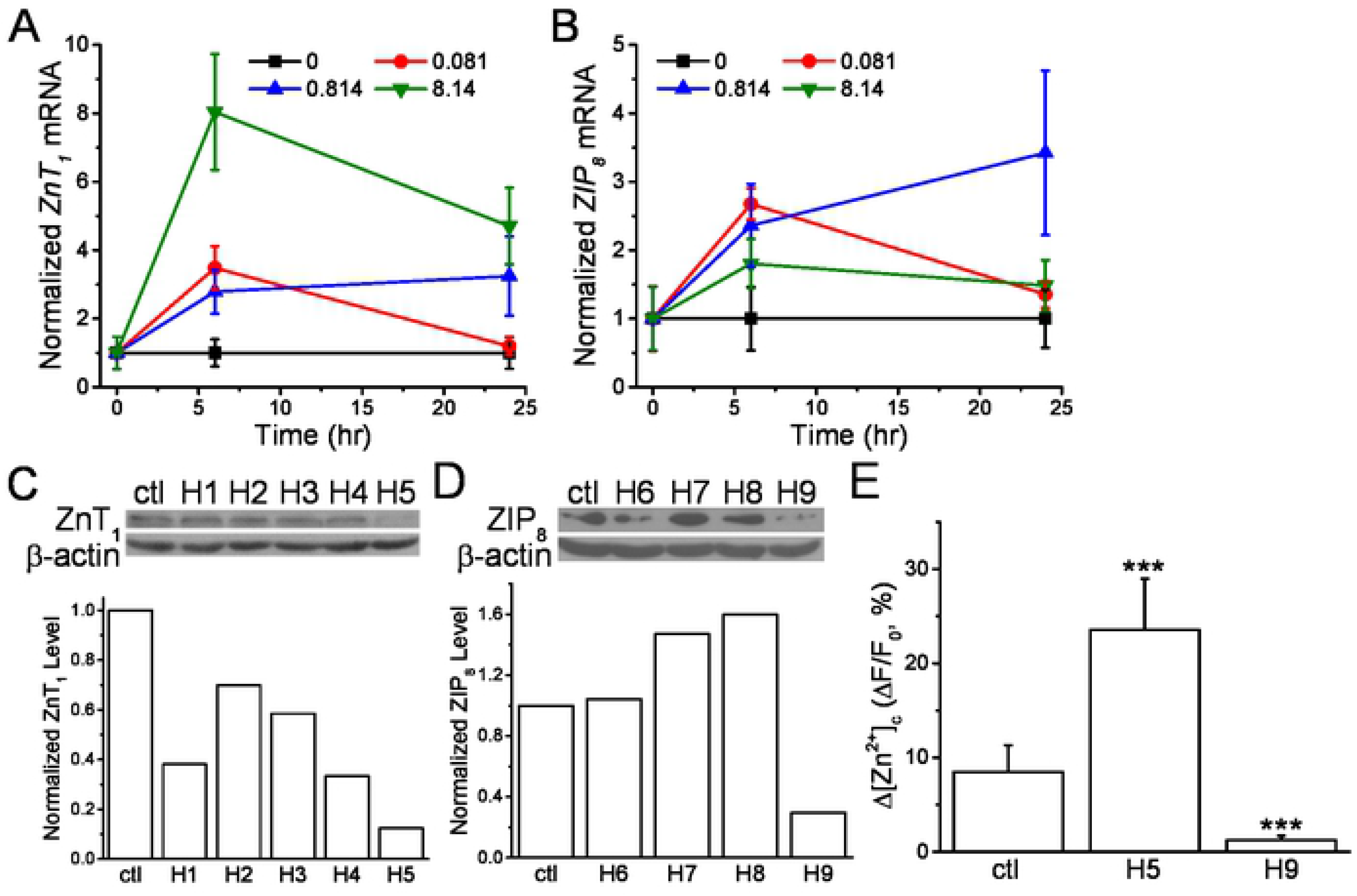
Knockdown the expressions of specific Zn^2+^ transporters interfere [Zn^2+^]_c_ responses in SH-SY5Y cells. A. and B. The expression levels of *ZnT1* and *ZIP8*, respectively. Cells were treated with different concentrations of ZnO-NP for 0, 6 and 24 hr and the mRNA levels of *ZnT*_*1*_ and *ZIP*_*8*_ were analyzed by RT-PCR. The expression levels were normalized to that of β-actin. C. and D. Expression knockdown of *ZnT1* and *ZIP8*, respectively. Specific shRNAs against *ZnT*_*1*_ (H1-5) and *ZIP*_*8*_ (H6-9) were delivered into the cells for 1 day and the protein levels were examined by Western blot (upper panel). The intensities of each protein bands were normalized to that of β-actin (lower panel). E. [Zn^2+^]_c_ responses in transfected cells. Cells were transfected with H5 and H9 shRNAs for 1 day and then loaded with FluoZin3. The changes in the fluorescence intensities (ΔF/F_0_) induced by ZnO-NP (0.814 μg/ml) were calculated. Data presented were Mean ± S.E.M from 3 bathes of cells. ***: *p* < 0.001 (Student’s *t*-test) when compared to the control group.

To verify the contributions of these transporters in regulating the [Zn^2+^]_c_ responses induced by ZnO-NP, we delivered specific shRNAs into the cells to reduce the translation of *ZnT*_*1*_ and *ZIP*_*8*_ (Fig. 2C & D, respectively). The results of the Western blots revealed that most of these shRNAs decreased the protein levels of *ZnT*_*1*_ (H1-5) and *ZIP*_*8*_ (H6-9); among them, H5 and H9 were the most effective shRNAs in reducing the protein levels of *ZnT1*, by 88%, and *ZIP*_*8*_, by 70%, respectively. Treating transfected SH-SY5Y cells with ZnO-NP (0.814 μg/ml), the averaged changes in [Zn^2+^]_c_, comparing to the control group, was about 4-fold higher in cells expressing H5 and mostly abolished in cells expressing H9 (Fig. 2E). It is likely that cells change the expression levels of these transporters to regulate the [Zn^2+^]_c_ in response to different stimulations.

### 3. ZnO-NP at a low dose increases the *Bax*/*Bcl-2* expression level

To characterize the toxicity of ZnO-NP on SH-SY5Y cells, we treated the cells with different concentrations of ZnO-NP for 24 hr and monitored the viability by MTT assay (Fig. 3A). The results show that ZnO-NP exposure reduced the viability in a dose dependent manner with an EC_50_ of 6.8 ± 0.2 μg/ml. Under 2 μg/ml, ZnO-NP had little effect on cell viability. We then examined the expression levels of *Bax* and *Bcl-2* by qPCR in SH-SY5Y cells treated with ZnO-NP at 0.081 and 0.814 μg/ml for 6 hr (Fig. 3C). The amounts of the PCR products expressed from *Bax* and *Bcl-2* decreased and increased, respectively, as the concentrations of ZnO-NP increased; in contrast, ZnO-NP at 8.14 μg/ml significantly increased the ratio to 1.49 ± 0.2. Therefore, that ZnO-NP at low non-lethal dose decreases the *Bax/Bcl-2* ratio indicating the blockage of apoptosis pathway.

**Figure 3.**
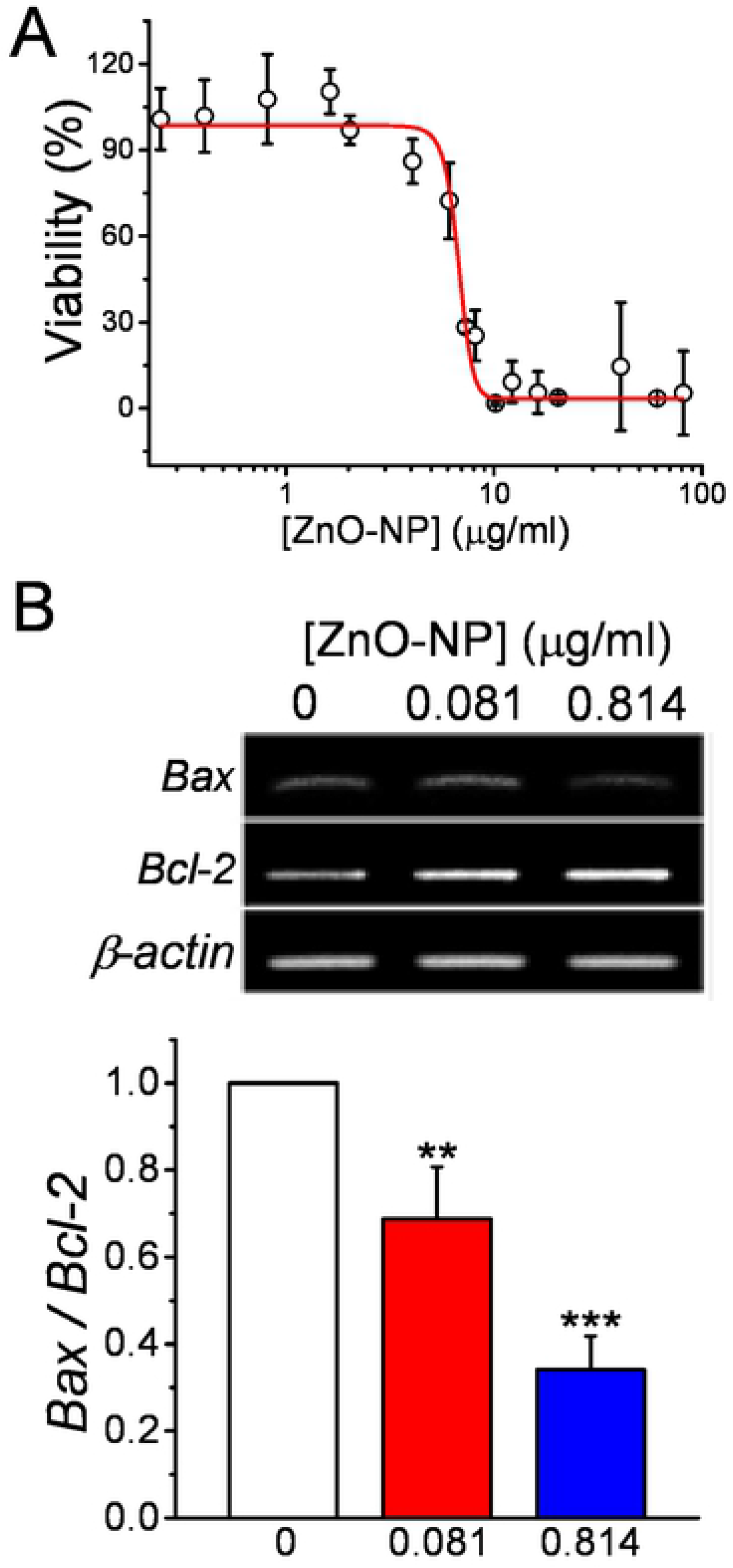
Low-dose ZnO-NP exposure reduces basal apoptosis signal in SH-SY5Y cells. A. Dose-dependent cell viability. After a 24-hr ZnO-NP exposure at different concentrations, the cell viability was analyzed by an MTT assay. The dose-dependence were fitted by a Boltzmann equation with an EC_50_ of 6.8 ± 0.2 μg/ml. Data presented were Mean ± SEM from 15 batches of cells. B. The *Bax/Bcl-2* ratio. Cells were treated with ZnO-NP for 6 hr and then the mRNA were collected for RT-PCR to analyze the expression level (upper panel) of *Bax* and *Bcl-2*. The intensities of the PCR products were normalized to the level β-actin and then used to calculate the *Bax*/*Bcl-2* ratio (Lower panel). Data presented were Mean ± S.E.M from 3 bathes of cells. ** and ***: *p* < 0.01 and 0.001, respectively, by Student’s *t*-test when compared to the control group.

ROS accumulation can trigger the expression of apoptosis-related genes. We then examined the intracellular ROS levels by loading the cells with H_2_DCFDA and monitored the changes in the fluorescence intensities in 2 hr (Supplementary Figure S2). For control cells without ZnO-NP treatment, the ROS level increased over the recording period; in the presence of ZnO-NP (0.081 and 0.814 μg/ml), the ROS levels at the same duration were lower than that of control group. These findings suggest that a low-dose exposure of ZnO-NP elicits beneficial effects in cells to reduce the oxidation stress and protect cells from death. b

### 4. ZnO-NP counteracted stress-induced ROS generation and cell death in SH-SY5Y cells

The uptake of 6-OHDA, an analog of dopamine, into cells through dopamine transporters triggers the production of ROS and causes cell death. To verify ZnO-NP has a protective effect on the 6-OHDA-induced cell death, we pretreated the SH-SY5Y cells with a low dose of ZnO-NP (0.081 and 0.814 μg/ml), which showed little effect on cell death in 24 hr (Fig. 3A). We then added 6-OHDA and monitored the survival rate at 6 hr later (Fig. 4). The results show that 6-OHDA significantly caused cell death with a dose-dependent manner at 50 and 100 μM. ZnO-NP pretreatment counteracted the 6-OHDA-induced cell death and became significant at 100 μM of 6-OHDA. In addition, ZnO-NP (0.081 μg/ml) pretreatment significantly suppresses the 6-OHDA-induced production of ROS. Similarly, ZnO-NP pretreatment reversed the effects of H_2_O_2_ in cell survival and ROS production (Supplementary Figure S3). We then used RT-PCR to examine the expression level of *p53*, a transcription factor involved in the activation of apoptosis pathway, in SH-SY5Y cells (Fig. 4C). It is apparently that ZnO-NP pretreatment reduced the expression of *p53* enhanced by 6-OHDA. These results suggest that ZnO-NP at a concentration below 1 μg/ml suppresses the production of ROS and facilitates cell survival.

**Figure 4.**
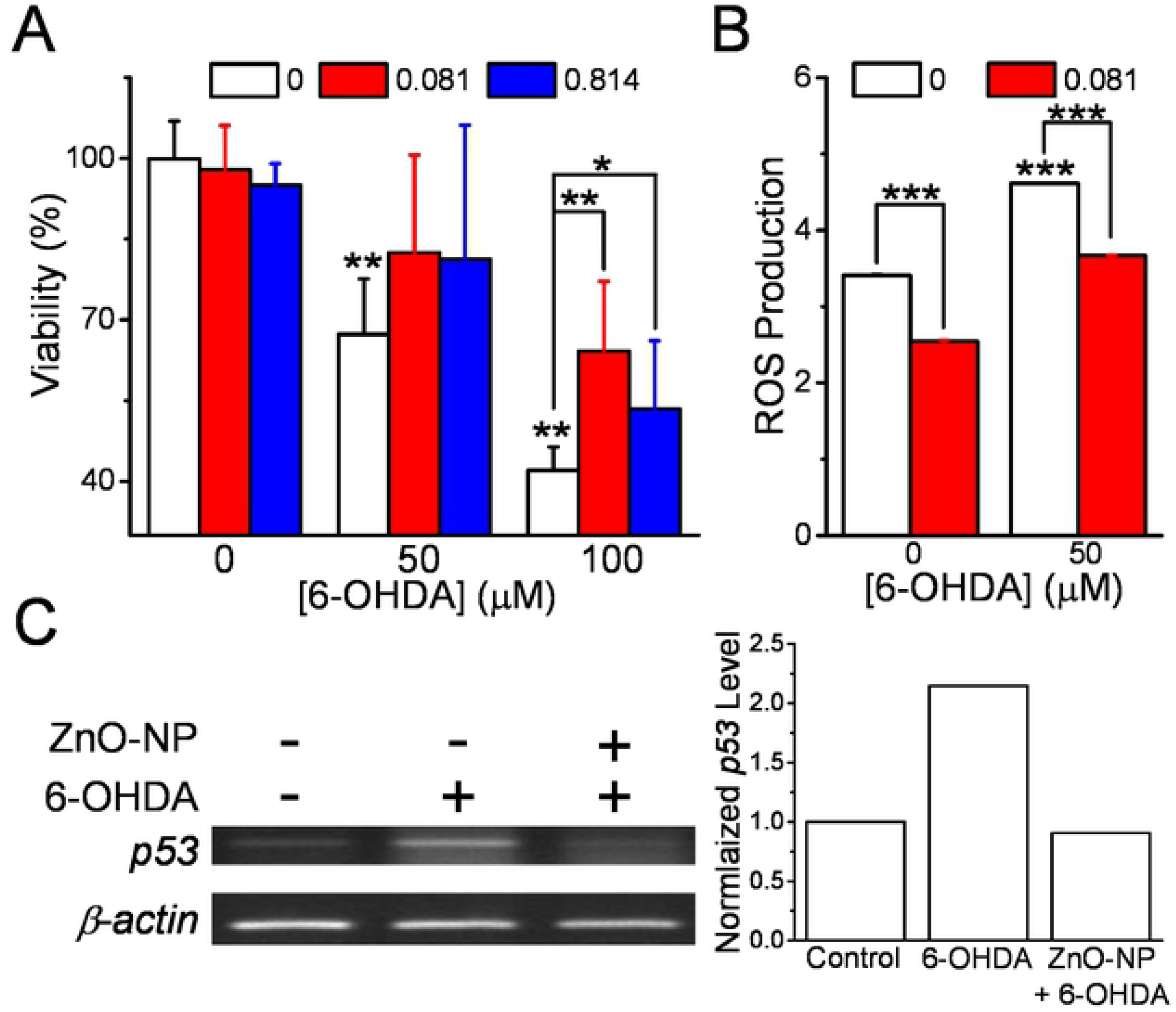
ZnO-NP suppresses 6-OHDA-induced cytotoxicity in SH-SY5Y cells. A. Cell viability. Cells were pretreated with ZnO-NP (0, 0.081, and 0.814 μg/ml) for 18 hr and then incubated with 6-OHDA (50 or 100 μM) for another 6 hr. The viability was measured by an MTT assay. B. ROS production. Cells were pretreated with or without ZnO-NP (0.081 μg/ml) for 2 hr, then 6-OHDA were added for another 1 hr. Data presented were Mean ± SEM from 15 batches of cells. The significance were analyzed by Student’s *t*-test; ** and ***: *p* < 0.01 and 0.001, respectively, when compared to the control group without 6-OHDA treatment or as indicated. C. *p53* mRNA levels. Cells were pretreated with ZnO-NP (0.081 μg/ml) for 18 hr and then incubated with 6-OHDA for 6 hr. Cells were then harvested for RT-PCR and the density of *p53* products were normalized with that of β-actin and control group.

### 5. ZnT1 and ZIP8 knockdown affected 6-OHDA-induced cytotoxicity

To verify the importance of ZnO-NP-induced elevation of [Zn^2+^]_c_ in protecting cells from death, we transfected the SH-SY5Y with shRNAs against *ZnT*_*1*_ and *ZIP*_*8*_, then examined the cell viability under 6-OHDA treatment with MTT assay (Fig. 5). The results show that knockdown the expression of *ZnT1* recused the cell death caused by 6-OHDA to a level similar to that of the control group and the addition of ZnO-NP did not enhance any more. In contrast, *ZIP*_*8*_ knockdown did not have such a protective effect in 6-OHDA-induced cell death and the addition of ZnO-NP did not reverse the toxic effect of 6-OHDA. As shown in Fig. 2E, knockdown the expression of *ZnT*_*1*_ and *ZIP*_*8*_ enhanced and suppressed the ZnO-NP-induced elevations of [Zn^2+^]_c_, respectively. Therefore, the release of Zn^2+^ from the acidic compartments by ZIP_8_ and the elevation of [Zn^2+^]_c_ facilitated by ZnT_1_ are important in enhancing the viability of cells under different challenges.

**Figure 5.**
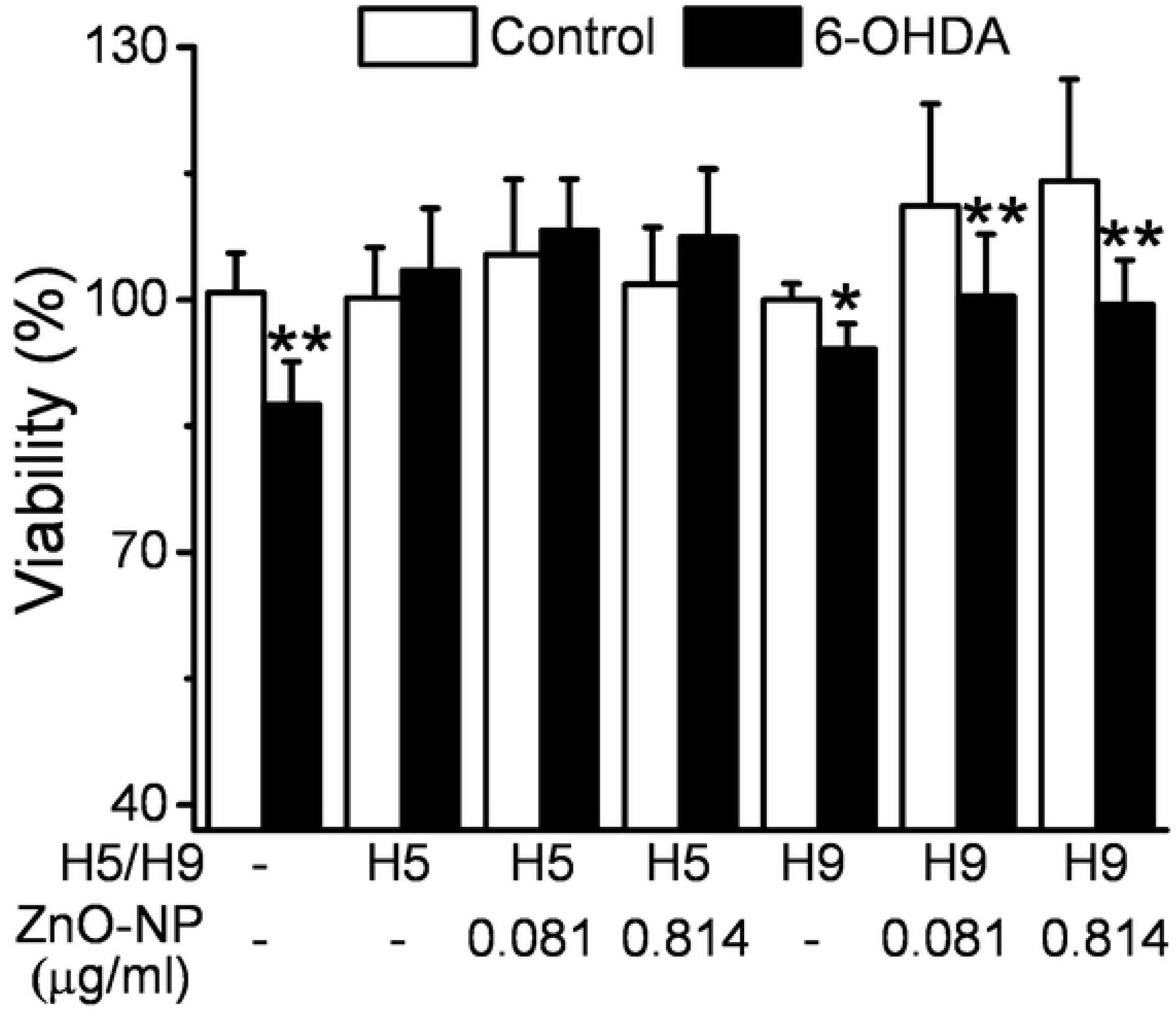
ZnO-NP altered 6-OHDA-induced cytotoxicity in cells with transporter knockdown. H5 and H9 snRNAs were transfected into SH-SY5Y cells for 24 hr to knock down ZnT_1_ and ZIP_8_, respectively. Cells were pretreated with ZnO-NP of different concentrations for 16 hr and then 6-OHDA (50 μM) for another 6 hr. The cell viability was determined by MTT assay. Data presented were Mean ± SEM (n = 15) and the significance were analyzed by Student’s *t*-test; * and **: *p* < 0.05 and 0.01, respectively, when compared to the group without 6-OHDA treatment.

## Discussion

This study finds that ZnO-NP potently induced the expressions of *ZnT*_*1*_ and *ZIP*_*8*_ to modulate [Zn^2+^]_c_, a crucial parameter for cytoviability in human neuroblastoma SH-SY5Y cells. Below lethal dosage under 1 μg/ml, ZnO-NP transiently elevated the [Zn^2+^]_c_ and decreased the *Bax/Bcl-2* expression ratio. In addition, ZnO-NP suppressed the cytotoxicity, ROS production and *p53* gene expression induced by 6-OHDA or H_2_O_2_. These results suggest the cell-protective function of ZnO-NP at lose dosages against oxidative stresses and support a therapeutic strategy by delivering ZnO-NP into the CNS to suppress the development of neuropathological disorders.

Zn^2+^ trafficking was investigated in these experiments. ZnO-NP-induced [Zn^2+^]_c_ changes were studied in cells transfected with snRNA against *ZnT*_*1*_ to illustrate the role of ZnT_1_ for the efflux of Zn^2+^. [Zn^2+^]_c_ and the expression of *ZnT*_*1*_ were coupled; both showed increases under exposure to low dose ZnO-NP and returned to the basal levels after 24 hr. At high dosage (8.14 μg/ml), ZnO-NP induced a large increase in [Zn^2+^]_c_, coupled with an 8-fold increase in *ZnT*_*1*_ mRNA (at 6 hr). In this case, both the expression level of *ZnT*_*1*_ and [Zn^2+^]_c_ remained high throughout the observation period. Moreover, neurotoxicity induced by 6-OHDA was suppressed in the *ZnT1*-kockdowned cells. Our data show that [Zn^2+^]_c_ changes are coupled with the *ZnT*_*1*_ expression levels which are closely related to the neuron-protection activity of Zn^2+^. ZnT_1_ is known to be a plasma membrane protein that is enriched in postsynaptic dendritic spines and plays a role in Zn^2+^ homeostasis in synaptic neuron functions and diseases (29). Su *et al.* reported a positive correlation between ZnT_1_ and Zn^2+^ content in the spinal cord (30), and ZnT_1_ is shown to increase significantly with progression of Alzheimer’s disease (31).

Our data suggest that changes in *ZnT*_*1*_ expression can become a marker for [Zn^2+^]_c_ disturbance associated with neuroviability. Other ZnTs such as ZnT_10_, at Golgi, is down-regulated by an elevation of extracellular Zn^2+^ in SH-SY5Y cells (32). IL-6 induces a down-regulation of *ZnT*_*10*_ and enhances the accumulation of Mn^2+^ that might be correlated with Parkinson’s disease (33). Further studies on ZnTs, ZIPs, and metallothioneins (MTs), are required to understand their roles in modulating the Zn^2+^ homeostasis.

We have previously demonstrated the internalization of ZnO-NP by PC12 cells upon exposure to the nanoparticles for 10 min. Furthermore, after nasal exposure to airborne ZnO-NP, the nanoparticles are found in rat brain under a transmission electron microscope (20). We also verify that ZnO-NP elevates [Zn^2+^]_c_ in both cultured cells and rat white blood cells through endocytosis and subsequent dissolution in acidic compartments such as endosomes (21). Conversion of ZnO to ions following entrance into lysosomes has also been shown in the studies of Xia *et al.* in which the labeled ZnO was traced in BEAS-2B cells (34). Muller *et al.* also have demonstrated that ZnO dissolves rapidly in a lysosomal fluid at a pH of 5.2 (35).

ZIP_8_ has been shown to be localized in the lysosomal membrane or in synaptosomes (27, 28). Our data show that ZnO-NP-induced [Zn^2+^]_c_ changes are greatly suppressed in *ZIP*_*8*_-knockdowned cells, illustrating that *ZIP*_*8*_ is required for intracellular Zn^2+^ release from those organelles after ZnO-NP was engulfed, which may be the main route for ZnO in elevating [Zn^2+^]_c_. The mRNA levels of *ZIP*_*8*_ and *ZnT*_*1*_ were positively correlated with the changes in [Zn^2+^]_c_ under exposure to ZnO-NP below 1 μg/ml. At a high dose of ZnO-NP (8.14 μg/ml), the expression of *ZIP*_*8*_ was small in contrast to [Zn^2+^]_c_ response and *ZnT*_*1*_ expression. The low level of ZIP_8_ prevent additional Zn^2+^ fluxing to the cytosol and further cellular damage. These results suggest that there is a negative feedback between elevation of [Zn^2+^]_c_ and the expression of *ZIP*_*8*_.

ROS is known to cause DNA damage that activates the *p53*-linked apoptosis pathway through phosphorylation by ATM. *Bcl-2* has been shown to be coupled with the pro-survival pathway to counteract effects of mitochondrial damage induced by *Bax*. In addition, silencing the expression of *ZnT*_*1*_, but not *ZIP*_*8*_, can not only enhance the ZnO-NP-induced [Zn^2+^]_c_ elevation but rescue the 6-OHDA-induced cell death. It is likely that [Zn^2+^]_c_ response is a perquisite for ZnO-NP to reduce stress-induced cytotoxicity by suppressing ROS generation and augmenting expression of *bcl-2*.

Zn^2+^ has been widely shown as a potential antioxidant for suppression of apoptosis (36–43). In animal brain studies, Zn^2+^ treatment decreases the *Bax*/*Bcl-2* protein ratio (43); treating SH-SY5Y cells with a low dose of Zn^2+^ can reverse a stress-induced increment of DNA fragmentation (12). Zn^2+^ deficiency has been shown to reduce stem cell proliferation, increase neuronal precursor apoptosis and impair neuronal differentiation (15, 44) as well as associate with neuronal dysfunction, such as attention-deficit hyperactivity disorder (9) and depression-like symptoms (45, 46). In contrast, Zn^2+^ supplementation can reduce the levels of ROS to prevent cardiomyocyte apoptosis and congenital heart defects (39); it also promotes the recovery of spinal cord function (17, 47). Zn^2+^ has a protective effect on renal ischemia-reperfusion injury by augmenting superoxide dismutase activity and lowering the *Bax/Bcl-2* expression ratio to reduce apoptosis (36). Our results support that ZnO-NP, at sub-lethal dosage, plays a protective role by reducing ROS generation and the expression of *Bax*/*Bcl-2*.

In this and previous studies, we show that ZnO-NP dose-dependently exert paradoxical protective and cytotoxic functions through their ability to alter [Zn^2+^]_c_ and modulate the expression of *ZnT*_*1*_ and *ZIP*_*8*_. Delivering ZnO-NP at a low dose into the central nervous system may provide a practical strategy to elevate the [Zn^2+^]_c_ for potent neuroprotection. Further studies, both *in vivo* and *in vitro,* will be required using more sensitive and selective techniques to measure the homeostasis of [Zn^2+^]_c_ and to assess the feasibility of using ZnO-NP for clinical application.

## Acknowledgements

We wish to thank Ms. Suzanne Hosier for English editing and Hui-Hsing Hung for the [Zn^2+^]_c_ measurements in normal neuronal cells. This work was supported by grants from Ministry of Science and Technology, Taiwan, R. O. C. (PSL: NSC 102-2320-B-031-001, MOST 104-2622-B-031-001-CC2, & MOST 104-2632-B-031-001; CYP: MOST 107-2320-B-002-052)

